# Subject-Agnostic Transformer-Based Neural Speech Decoding from Surface and Depth Electrode Signals

**DOI:** 10.1101/2024.03.11.584533

**Authors:** Junbo Chen, Xupeng Chen, Ran Wang, Chenqian Le, Amirhossein Khalilian-Gourtani, Erika Jensen, Patricia Dugan, Werner Doyle, Orrin Devinsky, Daniel Friedman, Adeen Flinker, Yao Wang

## Abstract

**Objective:** This study investigates speech decoding from neural signals captured by intracranial electrodes. Most prior works can only work with electrodes on a 2D grid (i.e., Electrocorticographic or ECoG array) and data from a single patient. We aim to design a deep-learning model architecture that can accommodate both surface (ECoG) and depth (stereotactic EEG or sEEG) electrodes. The architecture should allow training on data from multiple participants with large variability in electrode placements and the trained model should perform well on participants unseen during training.

**Approach:** We propose a novel transformer-based model architecture named SwinTW that can work with arbitrarily positioned electrodes by leveraging their 3D locations on the cortex rather than their positions on a 2D grid. We train subject-specific models using data from a single participant and multi-patient models exploiting data from multiple participants.

**Main Results:** The subject-specific models using only low-density 8×8 ECoG data achieved high decoding Pearson Correlation Coefficient with ground truth spectrogram (PCC=0.817), over N=43 participants, outperforming our prior convolutional ResNet model and the 3D Swin transformer model. Incorporating additional strip, depth, and grid electrodes available in each participant (N=39) led to further improvement (PCC=0.838). For participants with only sEEG electrodes (N=9), subject-specific models still enjoy comparable performance with an average PCC=0.798. The multi-subject models achieved high performance on unseen participants, with an average PCC=0.765 in leave-one-out cross-validation.

**Significance:** The proposed SwinTW decoder enables future speech neuropros-theses to utilize any electrode placement that is clinically optimal or feasible for a particular participant, including using only depth electrodes, which are more routinely implanted in chronic neurosurgical procedures. Importantly, the generalizability of the multi-patient models suggests that such a model can be applied to new patients that do not have paired acoustic and neural data, providing an advance in neuroprostheses for people with speech disability, where acoustic-neural training data is not feasible.

## 1. Introduction

Brain-related speech disability, which can be caused by stroke, injury, or tumor [10, 35, 44], can seriously decrease a patient’s quality of life. In the United States, an estimated 2.5 million people suffer from speech disability due to stroke alone [20]. There has been growing interest in using intracranial electrodes to record neural activity during speech production in order to directly decode human speech from these signals, making it possible to design Brain-Computer Interface to allow patients with speech disabilities to communicate [38, 30, 7, 33, 8, 36].

Recent advancements in deep neural networks have been leveraged to push the boundary of speech decoding from ECoG signals. The decoding pipeline proposed in [47, 9] first applies a Neural Decoder (called ECoG Decoder) to predict time-varying speech parameters and then uses a novel Speech Synthesizer to generate speech spectrograms from speech parameters. Using ResNet [15] or 3D Swin Transformer [29] as the Neural Decoder, high speech decoding performance in terms of PCC between the decoded and ground-truth spectrograms has been achieved. In [1], densely connected 3D Convolutional Neural Networks (CNN) were applied to decode speech from ECoG signals. Besides CNN and Transformer, recurrent neural networks (RNN) and long short-term memory (LSTM) networks have also been explored as Neural Decoders [31, 4, 34]. Some approaches produced naturalistic reconstruction leveraging wavenet vocoders [1], generative adversarial networks [46], and unit selection [16], but with limited accuracy. These studies demonstrate that deep neural networks can decode speech information from the complex neural activity recorded by the ECoG signals. A recent study in one patient with implanted high-density ECoG electrodes [32] was successful in decoding naturalistic speech with high word decoding accuracy by leveraging the quantized HuBERT features [18] as an intermediate representation space and a pre-trained speech synthesizer which converts the HuBERT features into speech. However, HuBERT features do not carry speaker-specific acoustic information and thus can only be used to generate a generic speaker’s voice, requiring a separate model to translate the generic voice to a specific patient’s voice.

The deep neural networks in previous speech decoding studies have architecture designs with several limitations. First, architectures that use spatial convolution among electrodes, e.g., [1, 9, 47], are only applicable to grid electrodes like an ECoG array and hence do not work with strip or depth electrodes. Vision transformers’ absolute position embeddings and relative positional bias are also based on the 2D or 3D grid index [12, 28, 27, 29] and hence are only applicable to grid electrodes [9, 24, 40]. On the other hand, the implantation of depth electrodes (stereotactic EEG or sEEG) has been a more popular neurosurgical approach which does not require the removal of a large skull portion with reports of fewer surgical complications [19, 43]. Further, the approach and electrodes employed in sEEG are similar to those used in Deep Brain Stimulation (DBS), which has demonstrated long-term electrode safety, suggesting the possibility of chronic sEEG for speech neuroprostheses [17]. Multiple sEEG depth probes may be implanted, which can assay a wide range of deeper structures and thus may provide additional information not available from the surface of the cortex. Therefore, decoding speech from sEEG signals would have significant clinical advantages.

Secondly, models that use fully connected computations among the electrodes, e.g., [4, 31, 39], can only be trained for a specific participant, as the weights learned depend on the actual locations of the electrodes in the brain. Further, electrode placement varies quite widely across patients, and fully connected architectures can not be trained effectively with data from multiple participants. Even convolutional [9, 47, 1] models or transformers [9, 47, 24, 40] that leverage grid indices for position embedding cannot generalize well to different participants because they do not specifically consider the locations of the electrodes on the brain. Therefore, studies to date have developed subject-specific models, which suffer from small data challenges as they cannot leverage signals from multiple subjects. More critically, such an approach requires collecting training data for each participant, limiting its practical applicability.

Our study proposes a novel transformer-based Neural Decoder that does not rely on a regular grid structure. We call it the Swin transformer with temporal windowing (SwinTW). Instead of relying on the grid index, the model leverages the anatomical location of electrodes in the standardized brain template to learn the attention between electrodes. The proposed Neural Decoder achieved superior performances than ResNet and 3D Swin Transformer across 43 participants, given the same grid electrodes, reported in [9]. The model demonstrated further performance increase by leveraging the off-grid electrodes that cannot be utilized in the previous studies. Importantly, the model demonstrated promising performance given sEEG electrodes only across 9 participants. Most significantly, the SwinTW model can be effectively trained with data from multiple participants, and the resulting model can generalize well to participants outside the training cohort.

## 2. Method

### 2.1. Speech Decoding Framework

Our neural decoding framework is trained by following a 2-step approach proposed in our previous study [9], shown in Figure 1. In the first step of Speech-to-Speech training, a Speech Encoder is used to extract speech parameters at every time frame (e.g., pitch, formant frequencies, loudness) from the input speech spectrogram, and a differentiable Speech Synthesizer is designed to reconstruct the spectrogram from the speech parameters. The Speech Encoder and the Speech Synthesizer are trained to match the reconstructed spectrogram with the ground truth. In the second step of Neural-to-Speech training, the Neural Decoder is trained to predict the time-varying speech parameters from neural signals using the speech parameters generated by the Speech Encoder as the guidance. The predicted speech parameters from the Neural Decoder are fed to the trained Speech Synthesizer from step 1 to generate the predicted speech spectrogram, which is then converted to the predicted speech waveform.

**Figure 1.**
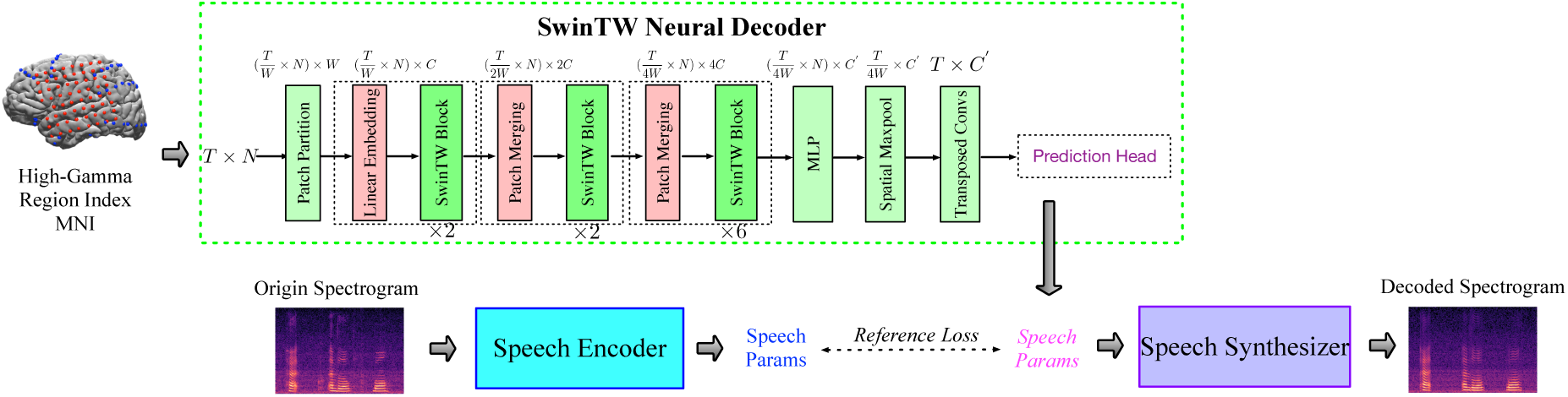
SwinTW Decoder Architecture and Speech Decoding Training Pipeline. SwinTW uses three stages of transformer blocks with spatial-temporal attention with temporal windowing to extract features. An MLP layer is applied to decrease the latent dimension after patch merging. After the last transformer block, spatial max pooling is applied across the electrodes to generate a single feature per time step. Finally, transposed temporal convolution is used to upsample the temporal dimension to be the same as the input. A prediction head module is used to generate speech parameters from the latent representation. The SwinTW is trained so that the decoded speech parameters from the neural signal match the reference speech parameters generated from the corresponding speech spectrogram using a pre-trained speech encoder. During inference, the speech encoder is not used. The SwinTW decodes the speech parameters from the neural signal, which are then fed into the speech synthesizer to generate the speech spectrogram.

Following the design from our previous study [47, 9], the Speech Encoder extracts 18 speech parameters at each time step from the original speech spectrogram, which is then fed to the Speech Synthesizer to reconstruct the original speech spectrogram. The Speech Encoder adopts a simple network architecture with MLP (Multilayer Perceptron) and temporal convolution. The differentiable speech synthesizer enables the end-to-end training of the speech-to-speech auto-encoding task (see bottom branch of Fig. 1). Details about the Speech Encoder and Speech Synthesizer and their pretraining using speech signal only can be found in [9].

For Neural-to-Speech training, the Neural Decoder first maps neural activity from all input electrodes to a latent feature, which is then used to predict the 18 speech parameters for each time frame, supervised by the speech parameters generated by the Speech Encoder. Then, the speech parameters predicted by the Neural Decoder will be fed into the Speech Synthesizer to generate the predicted spectrogram, which is then converted to the ECoG-decoded speech signal.

### 2.2. Neural Decoder based on Temporal Swin Transformer

In our study, we propose a novel architecture for decoding speech parameters from electrode signals that do not require electrodes to be on a 2D grid. We name the proposed Neural Decoder a Swin transformer with temporal windowing (SwinTW), inspired by the Swin Transformer [28, 27]. In the vanilla Vision Transformer (ViT) for an image [12], the self-attention layer computes global attention among all tokens (with each token corresponding to an image patch). This global attention causes the absence of the inductive bias of locality and heavy quadratic computational complexity to the input image size. The Swin Transformer solves the problems by grouping tokens into local windows and computing local attention within each window at each self-attention layer. To allow inter-window information exchange, the Swin Transformer shifts the window partition between every two windowed self-attention layers, which prevents different windows from being segregated (details can be found in [28, 27]). However, since the Swin Transformer was designed for 2D images (later extended to 3D videos [29]), its architecture assumes that the input is in the formats of 2D or 3D grids. Our previous transformer-based Neural Decoder used 3D Swin [9] which is inspired by [29], where each 3D window includes nearby 2 *×* 2 electrodes in two adjacent time steps. In our proposed SwinTW, we made several modifications to allow speech decoding based on electrodes in any topological layout. We still have spatial and temporal attention, but windowing is only applied in the temporal direction to constrain temporal attention. Instead of using electrode location on the 2D grid for spatial positioning information in [9], we use the anatomic location of each grid on the cortex (MNI coordinate and brain region index). The architecture of the SwinTW is shown in Figure 1.

#### Temporal patch partition

In the Swin Transformer [28, 27, 29] or ViT [12], the input images or videos are partitioned into 2D or 3D patches, and each patch is then mapped to a token with a patch embedding layer. This patch partition requires ordering all the electrodes into a 2D grid and makes the trained model not invariant to the electrode order. To solve this problem, our proposed SwinTW generates tokens from each electrode individually and only partitions the temporal dimension. As shown in Figure 1, given an ECoG signal with the shape of *T × N* (*T* : number of frames, *N* : number of electrodes), for each electrode, the SwinTW partitions the temporal sequence of neural activity into 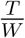 patches with patch size *W*. The temporal patch partition generates 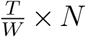 patches in total, and a linear patch embedding layer is applied to each patch to generate 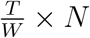 tokens with the latent dimension of *C*.

#### Temporal window attention

In Swin transformer [28, 27, 29], tokens are partitioned into windows, where each window contains a local subset of adjacent tokens, and attention is calculated only among tokens within the same window. In conventional 3D Swin, the windowing is applied spatially and temporally, making the model only suitable for electrodes arranged in a 2D grid. In SwinTW, to remove this grid input constraint, the model only partitions tokens into local windows in the temporal dimension and allows spatial attention across all electrodes (this can be thought of as using a spatial window size that includes all electrodes). Given *N* = *N*_*t*_ *× N*_*s*_ tokens (N: total number of tokens, *N*_*t*_: number of tokens in the temporal dimension, *N*_*s*_: number of tokens in the spatial dimension, equal to number of electrodes) and window size *W*_*t*_, the *N* tokens are partitioned into 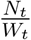 windows and attention is calculated among *W*_*t*_ *× N*_*s*_ tokens within each window.

#### Temporal patch merging

The Swin Transformer leverages patch merging to achieve inductive bias of locality and hierarchical feature maps. However, merging nearby patches in the spatial dimension is not feasible when the electrodes are not arranged in a grid. Therefore, instead of using the spatiotemporal patch merging in the 3D Swin Transformer [29], the SwinTW conducts temporal patch merging for each electrode individually. For each electrode, every two consecutive tokens in the temporal dimension with feature dimension *C* will be concatenated as a 2*C* dimensional latent and get mapped to a 2*C* dimensional merged token.

#### Grid-free positional embedding

The SwinTW follows Swin Transformers [27] to exploit positional information through relative positional bias. However, instead of using the 2D or 3D grid index difference as the relative position like the Swin Transformer, our SwinTW defines the relative positional bias based on each token’s anatomical location and time-frame index. The positional bias is defined as below:

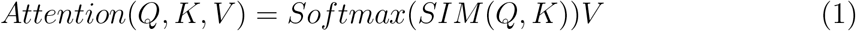

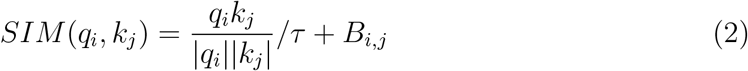

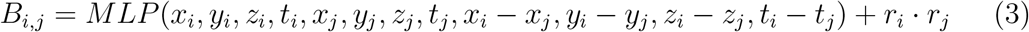

Given *Q, K, V ∈ R*^*N×C*^ (*Q, K, V* are query, key and value generated from each token, *N* is number of tokens and *C* is the latent dimension), shown in equation 1, the softmax of *SIM* (*Q, K*) for all pairs of token in the window is used to aggregated *V* (values of tokens within the window) to get the output token values. We define query-key similarity following the scaled cosine attention of SwinV2 [27], defined in equation 2. *τ* is a learnable parameter not shared among attention heads and layers. *B*_*i,j*_ is the relative positional bias between token *i* and token *j*. In SwinTW, *B*_*i,j*_ consists of two terms: MNI-based positional bias and region-based bias. We project each subject’s electrodes to a standardized Montreal Neurological Institute (MNI) brain anatomical map and collect each electrode’s *x, y, z* location in the MNI coordinate. For each token pair, the MNI coordinates of the corresponding electrodes and time-frame index, along with their differences, will be mapped to the MNI-based positional bias with a 2-layer MLP, which is shown in the first term of Eq. (3). We also parcellate the standardized brain into regions of interest (ROIs) and learn a dictionary of embeddings for all ROIs, with *r*_*i*_ denoting the embedding features for region *i*. Given *N*_*r*_ ROIs and *N*_*h*_ attention head, the learnable dictionary has *N*_*h*_ sets of *N*_*r*_ *×C*_*r*_ region embeddings (*C*_*r*_ is the region embedding dimension). The region embeddings *r*_*i*_, *∀i* are learned during the training. For a pair of tokens, the dot product of the embeddings of their corresponding electrodes’ ROIs will be added to the positional bias, shown in the second term of Eq. (3). The dot product is used instead of cosine similarity to allow the model to assign high attention to certain regions by letting them have large embedding values.

The architecture of SwinTW is shown in Figure 1. The input ECoG signal with a size of *T × N* is partitioned into 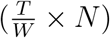 patches, each with a patch size of *W ×* 1. A linear patch embedding layer maps each patch to a *C* dimensional token. The SwinTW has three stages with 2, 2, and 6 layers. Swin Transformer Block (consists of a windowed multi-head self-attention layer and an MLP) is applied in each layer, detailed in [28], and we replace the spatial-temporal windowing with temporal-only windowing. Following the Swin Transformer, the temporal window partition is shifted for every two consecutive layers to allow inter-window information exchange, detailed in [28]. SwinTW performs temporal patch merging after the first and second stages, each stage decreasing the token number by half and doubling the latent dimension. After stage 3, an MLP is applied to decrease the 4*C* latent dimension to *C*^*′*^. Spatial max pooling across the electrodes is then applied to convert 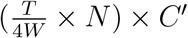 feature maps to 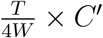 Transposed temporal convolutions are then employed to upsample 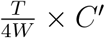 to 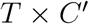, where *T* is the frame number of the input neural signal. As shown in 1, the *T × C*^*′*^ latent from SwinTW next goes through the prediction head consisting of temporal convolutions (kernel-size=3) and MLP to predict the 18 speech parameters at every frame.

In our study, we set *C* = 96 and *C*^*′*^ = 32. Patch-size *W* = 4 and window size *W*_*t*_ = 4 is applied to partition temporal dimension. In our 3 stages SwinTW with 2, 2, and 6 layers, the self-attention layers in the 3 stages have 3, 6, and 12 attention heads, respectively. The MLP in SwinTW has 3 layers (384→196→96→32) with layer norm [5] and LeakyRELU activation in between. The transposed convolution for temporal upsampling contains 4 1D transposed convolutional layers with stride=2 and kernel-size=3, padding=1. These parameter choices are determined through empirical trials and errors.

### 2.3. Training of Subject-Specific Neural Decoders

The training procedures for both the Speech Encoder and Speech Synthesizer follow the methods described in our previous work [9]. Therefore, we omit the details in this section. Following [9], we use two types of supervision to guide the training of the Neural Decoder that predicts speech parameters from neural signals. Firstly, we train the decoder to generate speech parameters that match the parameters generated by the speech encoder. Besides, the ground truth speech spectrograms act as additional supervision for the decoder, as the predicted speech parameters are converted to spectrograms by the speech synthesizer. The fact that our Speech Synthesizer is differentiable enables us to use the spectrogram reconstruction loss for end-to-end training. The reference loss *L*_*reference*_ for the speech parameters is defined as:

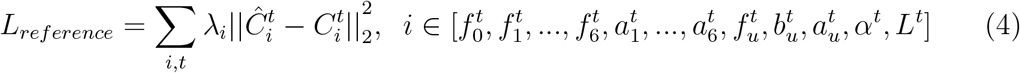

where 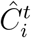 and 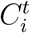 are speech parameters generated by the Neural Decoder and the Speech Encoder (as ground truth), respectively. We assign each speech parameter with individual weight *λ*_*i*_ through testing the performances on three hybrid-density participants with different parameter choices, and the values are detailed in [9]. For spectrogram-based supervision, we use modified multi-scale spectral loss *L*_*MSS*_, Short-Time Objective Intelligibility (STOI) loss *L*_*ST OI*_, and supervision loss *L*_*supervision*_. *L*_*MSS*_ is inspired by [13]. It supervises speech reconstruction by measuring the distance between the ground truth spectrogram and the reconstructed spectrogram in both linear and mel-frequency scales. *L*_*ST OI*_ measures the intelligibility of reconstructed speech based on the STOI+ metric [14]. Higher STOI+ indicates better intelligibility, the *L*_*ST OI*_ is defined as the negative of STOI+: *L*_*ST OI*_ = *−STOI*+. Besides, additional supervision *L*_*supervision*_ is applied to improve the prediction accuracy for the pitch 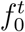 and formant frequencies 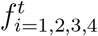 The *L*_*supervision*_ calculates the L2 distance between each predicted frequency and the corresponding frequency extracted by the Praat method [6]. The overall loss for training the Neural Decoder is

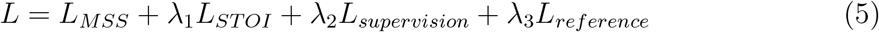

where *λ*_1_ *λ*_2_ and *λ*_3_ are set to 1.2, 0.1, and 1.0, following [9] through testing the performances on three hybrid-density participants with different parameter choices. Adam optimizer [22] with learning-rate=5 *×* 10^*−*4^, *β*_1_=0.9 and *β*_2_=0.999 is used to train the Neural Decoder. As mentioned in Section 3.1, following [9], randomly selected 50 out of 400 trials are used as the test set for each subject, and the remaining data are used for training.

### 2.4. Multi-Subject Neural Decoder Training

The proposed SwinTW allows the Neural Decoder to take input with any electrode layout as long as we know each electrode’s MNI coordinate and region index. Therefore, this architecture allows the Neural Decoder to be trained using data from multiple participants and then used for inference on any participant. Figure 2 demonstrates the multi-subject Neural Decoder training pipeline. Given data from multiple participants, a shared SwinTW-based Neural Decoder generates speech parameters based on each participant’s electrode signals and electrode locations (electrodes’ MNI coordinates and region index). Reference loss is calculated between the predicted speech parameters and the speech parameters generated by the subject-specific Speech Encoder. Each subject’s predicted speech parameters are fed into the corresponding subject-specific speech synthesizer to generate a speech spectrogram. The neural signals and electrodes’ locations are fed into the Neural Decoder to generate speech parameters during inference. The participant’s speech synthesizer then generates a speech spectrogram from the predicted speech parameters. Note that the embeddings for different ROIs are also learned as part of the Neural Decoder training. When we train a decoder using participants with left and right hemisphere electrodes, separate region embeddings are learned for the left and right brain hemispheres.

**Figure 2.**
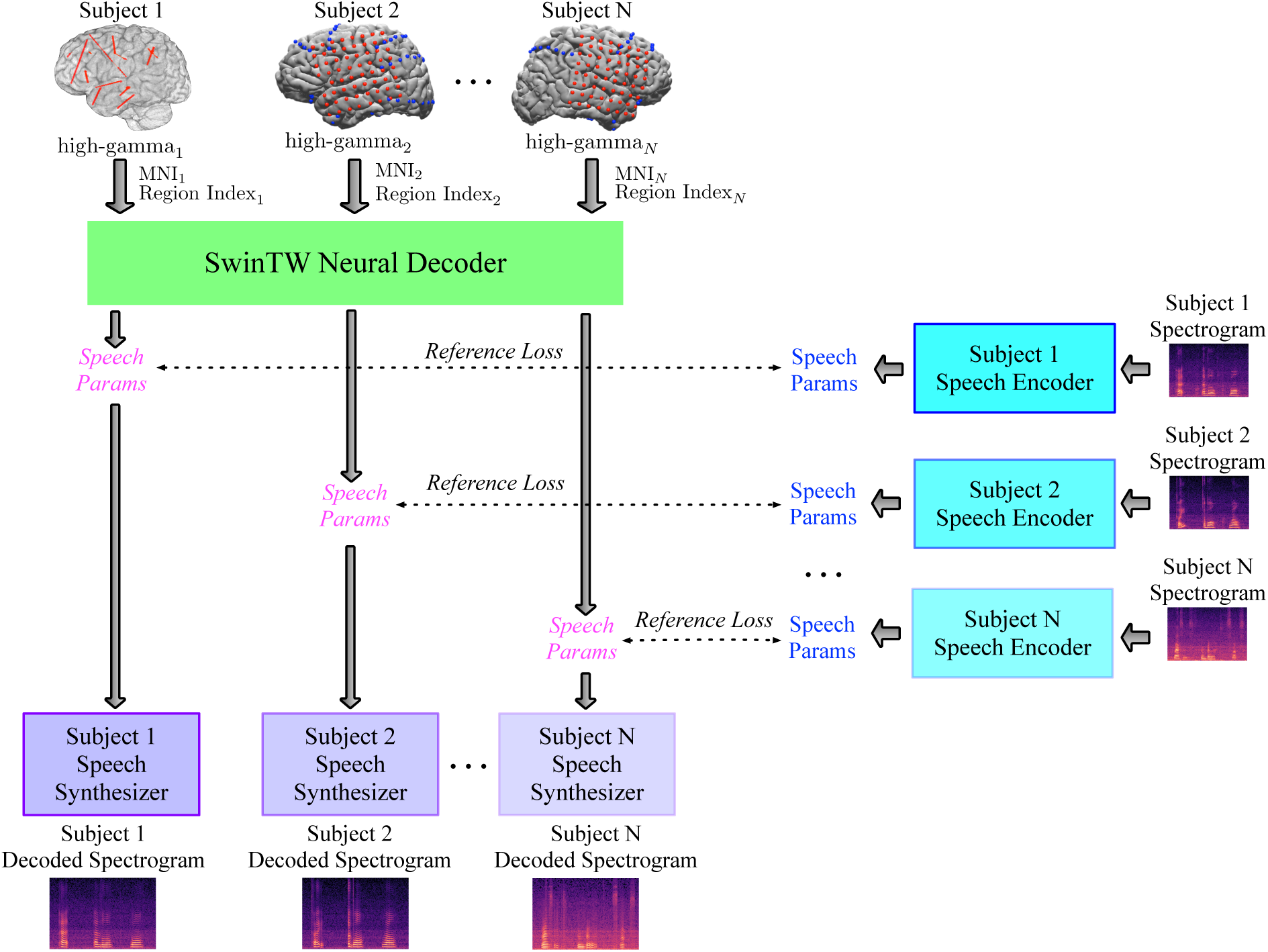
Multiple-subject Neural Decoder training pipeline. Each participant’s neural signal and electrodes’ location information (MNI coordinates and ROI index) are fed to a shared SwinTW Neural Decoder to predict speech parameters. The predicted speech parameters are supervised by the speech parameters generated by the subject-specific Speech Encoder from the ground-truth speech spectrogram. Each participant’s predicted speech parameters are fed into the corresponding subject-specific Speech Synthesizer to generate a speech spectrogram.

### 2.5. Evaluation Metrics

Following [48, 1, 4, 9], we used three metrics to evaluate the speech decoding performance:

Pearson Correlation Coefficient (PCC) measures the normalized correlation between the decoded spectrogram and the actual spectrogram and is a widely used metric to evaluate the accuracy of the decoded spectrogram.
STOI+ [14] is another metric that measures the similarity between decoded and original speech. STOI+ has been reported to have a monotonic relationship with speech intelligibility. The STOI+ value ranges from -1 to 1, and higher STOI+ indicates better intelligibility.
Mel-cepstral distortion (MCD)[25] measures the differences between 25 acoustic features generated from the decoded speech and the original speech. A lower MCD is better. MCD is calculated as follows:

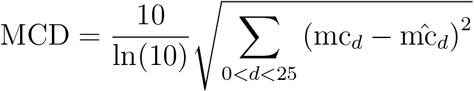

where mc_*d*_ and 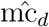 denote the *d*-th feature generated from the original and decoded speech.

## 3. Results

### 3.1. Neural Data Collection and Preprocessing

The study includes 52 native English-speaking subjects (43 subjects with ECoG electrodes, 20 males, 23 females; 9 subjects with only sEEG electrodes, 3 males, 6 females) with refractory epilepsy (a disease involving seizures caused by abnormal electrical brain activity). Details about speech and ECoG signals collection can be found in [9]. In brief, at each trial, a subject was requested to speak a specific target word in response to an audio or visual stimulus while their neural activity signals were recorded. Each subject was asked to complete 5 different tasks: (1) Auditory Repetition (repeating the word that the care provider has spoken), (2) Auditory Naming (naming the word based on the definition that the care provider has spoken), (3) Sentence Completion (naming the last word to complete a sentence that the care provider has spoken) (4) Visual Reading (reading the written word shown by the care provider) (5) Picture Naming (naming the word based on a colored drawing shown by the care provider). Each task included the same 50 target words [41], each appearing once in the Auditory Naming and Sentence Completion and twice in each of the other tasks, leading to 400 trials of ECoG signal recording, and the average duration of word production among all trials was 500ms.

All electrodes were implanted to capture clinically relevant brain regions, detailed in [9]. There were 43 subjects who had 8×8 ECoG electrodes with 10 mm spacing capturing signals over the perisylvian cortex (male left hemisphere: 14 subjects; female left hemisphere: 13 subjects; male right hemisphere: 6 subjects; female right hemisphere: 10 subjects). Besides the 8×8 grid electrodes, some subjects had additional electrode strips outside the 8×8 grid and/or depth electrodes implanted under the brain’s surface. We also included 9 subjects with only sEEG electrodes (male = 3, female = 6). The experiments were approved by the Institutional Review Board of NYU Grossman School of Medicine, and written and oral consent was collected from each participant. All implanted electrodes were the clinical standard of care and FDA-approved. The high gamma component (70-150 Hz) was extracted from the raw electrode signal, with electrodes exhibiting artifacts or interictal/epileptiform activity excluded by setting their signal to 0. The preprocessing details can be found in [9]. This study also applies a Savitzky-Golay filter [37] with a 3rd-order polynomial and window size of 11 to further denoise the high-gamma signal in the temporal dimension. Among the 400 trials of ECoG signals recorded from the five-word production tasks, 350 trials were used for model training, and 50 trials were held out for testing (10 randomly selected trials were reserved for testing for each task).

### 3.2. Subject-Specific Models: Speech Decoding with Electrodes on One ECoG Grid

To compare our proposed grid-free SwinTW with the Neural Decoders based on ResNet and 3D Swin transformer in our previous study [9], firstly, we evaluated the SwinTW trained with 64 ECoG electrodes for each subject individually. Figure 3 compares the decoding performance of the SwinTW decoder with the 3D ResNet and 3D Swin decoders. For each subject, we compute the average PCC, STOI+, and MCD among all the test trials. Each dot in a box plot is the mean metric for one subject. As illustrated in Figure 3, the SwinTW (PCC = 0.825, STOI+ = 0.309, MCD = 2.341) outperforms ResNet (PCC = 0.804, STOI+ = 0.264, MCD = 2.374) and 3D Swin transformers (PCC = 0.785, STOI+ = 0.216, MCD = 2.425) in terms of PCC, STOI+, and MCD. Additionally, the performance of the three models tested on shuffled data (by randomly shuffling the input neural signals temporally during the entire recording session) is also reported as a control in the supplementary Figure. S1. It is evident that the decoding performance with non-shuffled data is significantly better. Note that SwinTW differs from 3D Swin primarily in how the spatial positions of two electrodes affect the spatial attention bias between the two electrodes. With 3D Swin, the relative position between the two electrodes on the 2D grid determines the attention bias, whereas, with SwinTW, the attention bias depends on the MNI coordinates and ROI embeddings of these electrodes. Our results suggest that using the MNI coordinates and ROI information can lead to better decoding performance while making the model applicable to non-grid electrodes.

**Figure 3.**
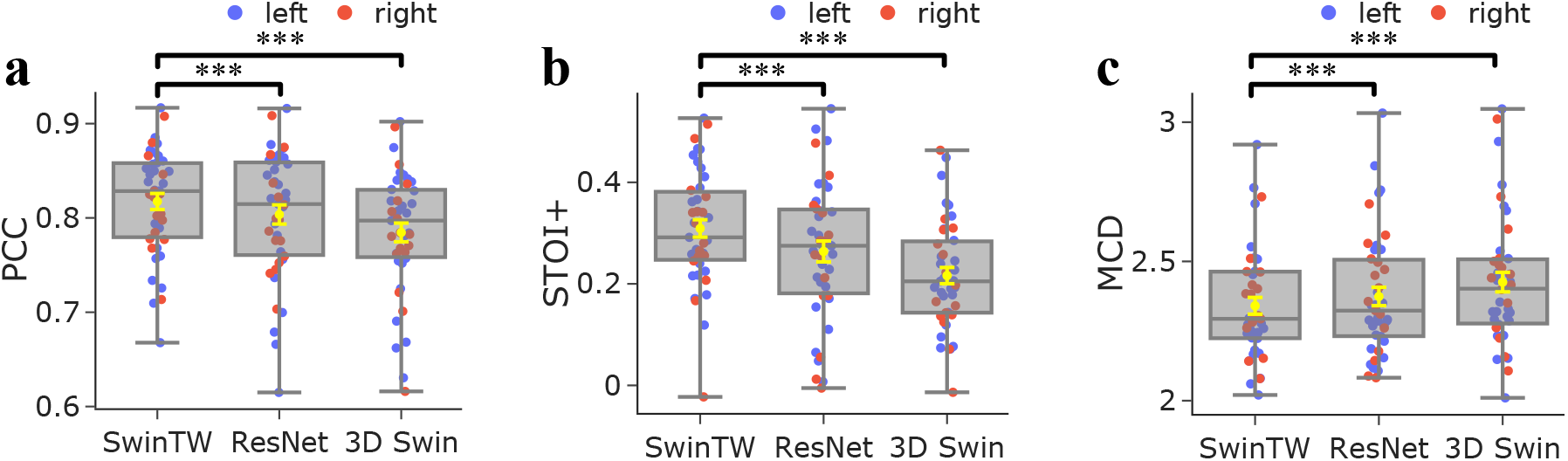
Performance comparison between different models. Subject-specific models when trained and tested on grid electrodes using different Neural Decoder architectures. Comparison of the distributions of decoding PCC **(a)**, STOI+ **(b)**, and MCD **(c)** over 43 participants. Each dot in a box plot indicates the mean metric for one participant across all testing trials. The yellow error bars denote the mean *±* standard error of the mean (SEM) across participants. The SwinTW outperforms the ResNet and 3D Swin Transformer regarding PCC and STOI+. All box plots depict the median (horizontal line inside box), 25th and 75th percentiles (box), 25th or 75th percentiles *±*1.5*×* interquartile range (whiskers) across all participants (N=43), and the yellow error bars denote the mean *±* standard error of the mean (SEM) across participants. Distributions were compared with each other as indicated. Black brackets indicate two experiments are compared using the Wilcoxon two-sided signed-rank test. ***: P *<* 0.001, *: P *<* 0.05, ns: p *>* 0.05.

### 3.3. Subject-Specific Models: Speech Decoding with Additional Electrodes

As the SwinTW does not rely on the 2D grid positions of the electrodes, the proposed SwinTW can easily leverage off-grid electrodes to provide additional information for speech decoding. In our study, for each participant with additional electrodes beyond one ECoG grid, we selected additional electrodes with a standard deviation of the signal greater than a subject-specific threshold, determined following the approach described in [21] for identifying active electrodes. We then trained the SwinTW Neural Decoder with 64 electrodes from the 8×8 grid and the selected additional electrodes for each subject. As 4 participants did not have any additional electrodes that fulfill the threshold requirement, we compared the models based on the remaining 39 participants. Each participant had 1 to 19 strip electrodes, 1 to 21 depth electrodes, 1 to 21 extra grid electrodes, and 1 to 11 electrodes with unknown locations (we set the MNI coordinates of these electrodes as 0 and the region index corresponding to Unknown instead of discarding them). Figure 4 compares the SwinTW Neural Decoder performance using all selected electrodes and with the performance using only electrodes on one ECoG grid.

**Figure 4.**
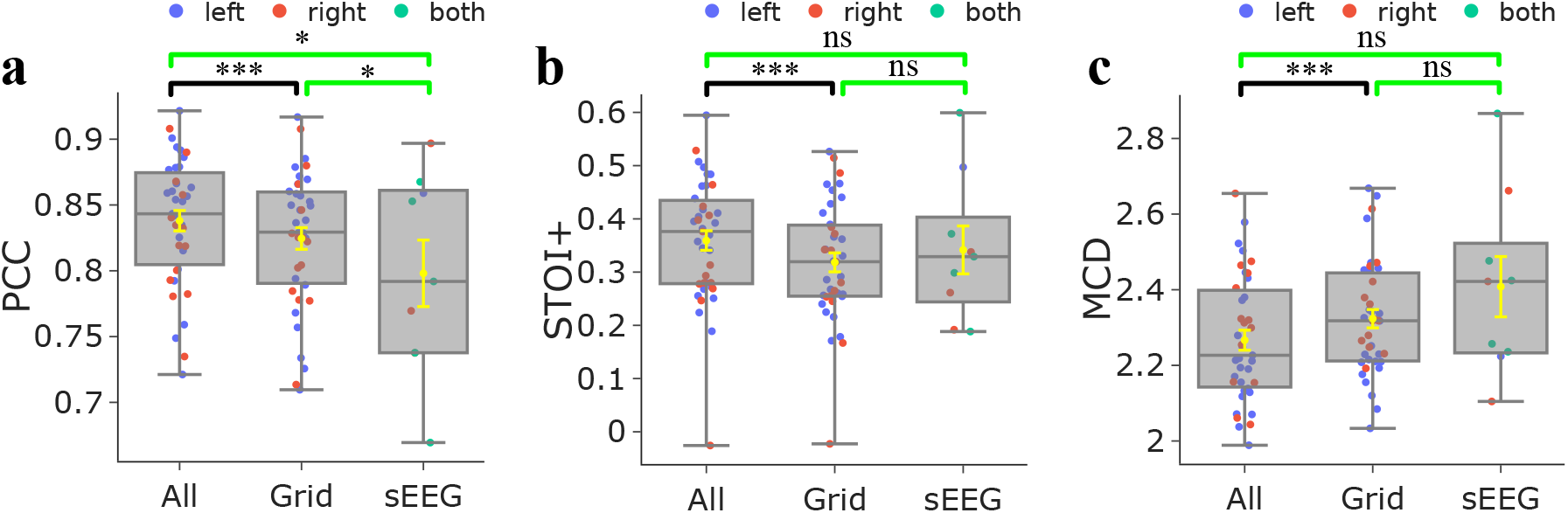
Performance comparison between different modalities with Subject-specific SwinTW model. Decoding PCC **(a)**, STOI+ **(b)**, and MCD **(c)** shows the comparison between subject-specific SwinTW Neural Decoder performance obtained with all selected electrodes, with only electrodes on one 8×8 grid for 39 participants, and with sEEG-only electrodes over 9 participants. Using all electrodes outperforms using grid electrodes or sEEG-only electrodes. Black brackets indicate two experiments are compared using the Wilcoxon two-sided signed-rank test. Green brackets indicate two experiments that are compared using the Wilcoxon rank-sum test, indicated in green. *∗ ∗ ∗* : *P* ¡0.001, *∗* : *P* ¡0.05, *ns* : *p*¿0.05.

The results demonstrate that additional electrodes can further improve the decoding performance (all electrodes PCC = 0.838, STOI+ = 0.359, MCD = 2.228; grid electrodes PCC = 0.825, STOI+ = 0.318, MCD = 2.341).

### 3.4. Subject-Specific Models: Speech Decoding with sEEG electrodes only

We also attempted to train the proposed SwinTW model to decode speech production from only SEEG electrodes. Our study included 9 subjects (male = 3, female = 6) with only sEEG electrodes implanted. For each subject, electrodes with a standard deviation of the signal greater than a subject-specific threshold derived following the approach of [21] were included. The number of selected electrodes for each participant ranges from 19 to 178. Figure 4 demonstrates that the SwinTW can achieve promising speech production prediction based on sEEG electrodes only, with the mean of PCC slightly lower (0.798 vs 0.825) but STOI+ (0.341) slightly higher than the decoding results from 43 participants with 64 ECoG grid electrodes (0.318).

### 3.5. Multi-Subject Model: Evaluation on Test Trials of Participants within the Training Set

As the proposed SwinTW architecture does not require the electrodes to be arranged in a grid but relies on the electrode position in the brain, it can handle the differences in the electrode placements among different participants and allow a single model to be trained with multiple patient data. To validate this idea, we trained a single SwinTW decoder with 15 randomly selected male participants with ECoG electrodes implanted in either the left or right brain hemisphere (4 on the left and 11 on the right). As detailed in Section 2.4, subject-specific speech encoder and speech synthesizer are applied while the Neural Decoder is shared among subjects. We compare the decoding performance of the multi-subject and subject-specific models on the test trials of each of the 15 participants included in the multi-subject model training. As shown in Figure 5, the multi-subject SwinTW model (PCC = 0.837, STOI+ = 0.352, MCD = 2.307) showed similar performance with the subject-specific model (PCC = 0.831, STOI+ = 0.334, MCD = 2.313), with no statistically significant differences.

**Figure 5.**
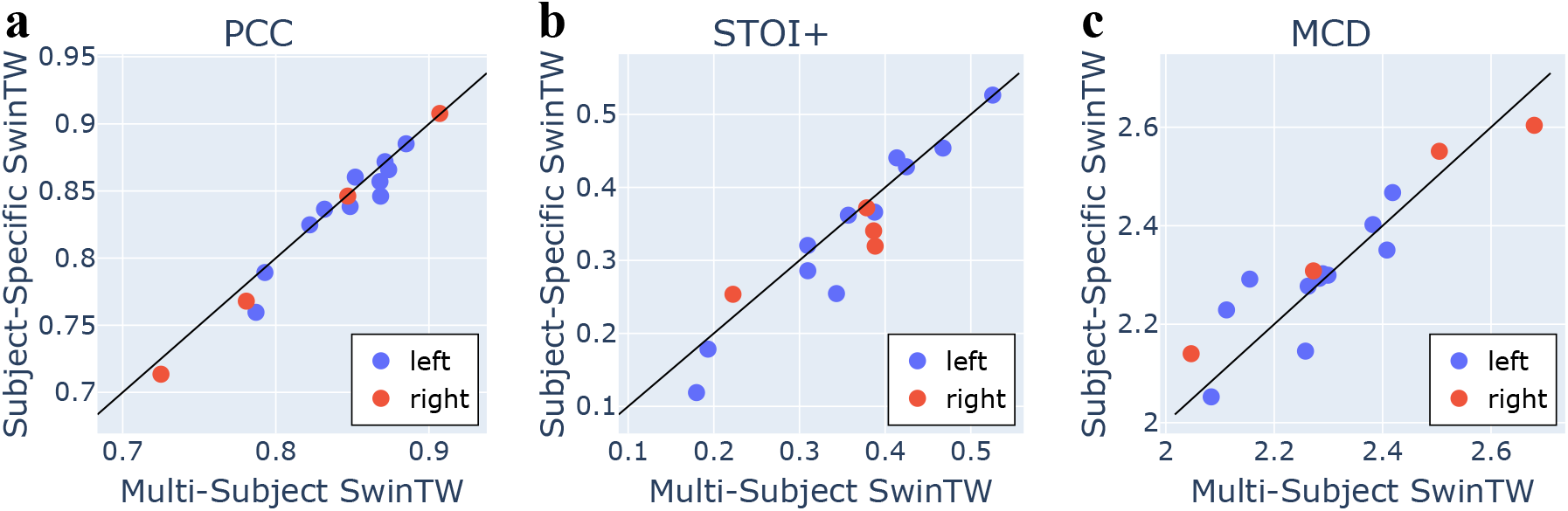
Comparison between the SwinTW Neural Decoder trained with 8×8 ECoG data from multiple (15) subjects and the subject-specific SwinTW. PCC, STOI+, and MCD were evaluated on test trials from the 15 participants. Wilcoxon’s two-sided signed-rank test is used to compare the two models. P-values of PCC, STOI, and MCD are 0.12, 0.06, 0.27.

### 3.6. Multi-Subject Model: Evaluation on Participants Outside the Training Set

We also evaluated the multi-subject SwinTW decoder on test trials of the subjects outside the training set. We conducted 5-fold cross-validation separately for male (n=20) and female (n=23) participants. Specifically, we partitioned all male (resp. female) participants (with ECoG electrodes implanted in either the left or right brain hemisphere) into five folds. Each time, we used data from four-fold participants to train a SwinTW decoder and evaluate its decoding performance on the remaining one-fold participants. The process is repeated to use every fold as the validation fold once. As shown in Figure 6, although the performance achieved by participants outside the training set is significantly lower than the subject-specific models, the decoded speech still has a high mean PCC of 0.765. The results demonstrate the proposed SwinTW decoder can achieve generalizability to participants unseen during model training.

**Figure 6.**
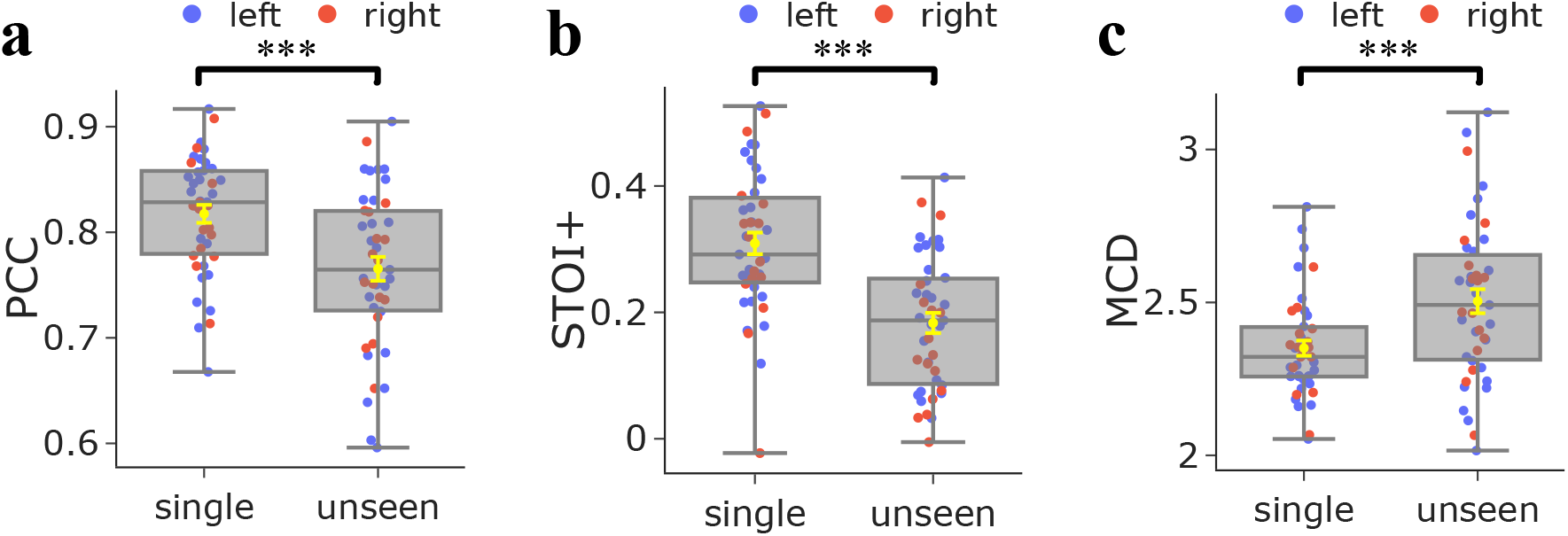
The decoding performance of the trained multi-subject model on participants outside the training set. Cross-validation was conducted on male and female subjects separately. Single refers to subject-specific model performance on each single subject, while unseen refers to the multi-subject model tested on unseen subjects. Although the performance is lower than subject-specific models, these results demonstrate that the SwinTW decoder trained with multi-subject data can generalize quite well to unseen subjects. Box plots as described in Fig. 3 across all participants (N=43), and the yellow error bars denote the mean *±* standard error of the mean (SEM) across participants. Distributions were compared with each other as indicated using the Wilcoxon two-sided signed-rank test for **a, b**, and **c**.***: P *<* 0.001.

To investigate if separate models should be trained for decoding from neural data in the left and right hemispheres, we performed additional experiments, where we trained and evaluated multi-subject models for the two hemispheres separately, each through cross-validation. Among male participants, there were 14 with left hemisphere data and 6 with right hemisphere data. For female participants, we had 13 with left hemisphere data and 10 with right hemisphere data. We used 5-fold cross-validation for training and evaluating each model. As shown in Figure 7, compared with hemisphere-specific models, the SwinTW decoder trained using data from both hemispheres achieved comparable performance when tested on unseen subjects. This suggests that a single SwinTW model can effectively extract and synthesize information from both hemispheres for speech decoding.

**Figure 7.**
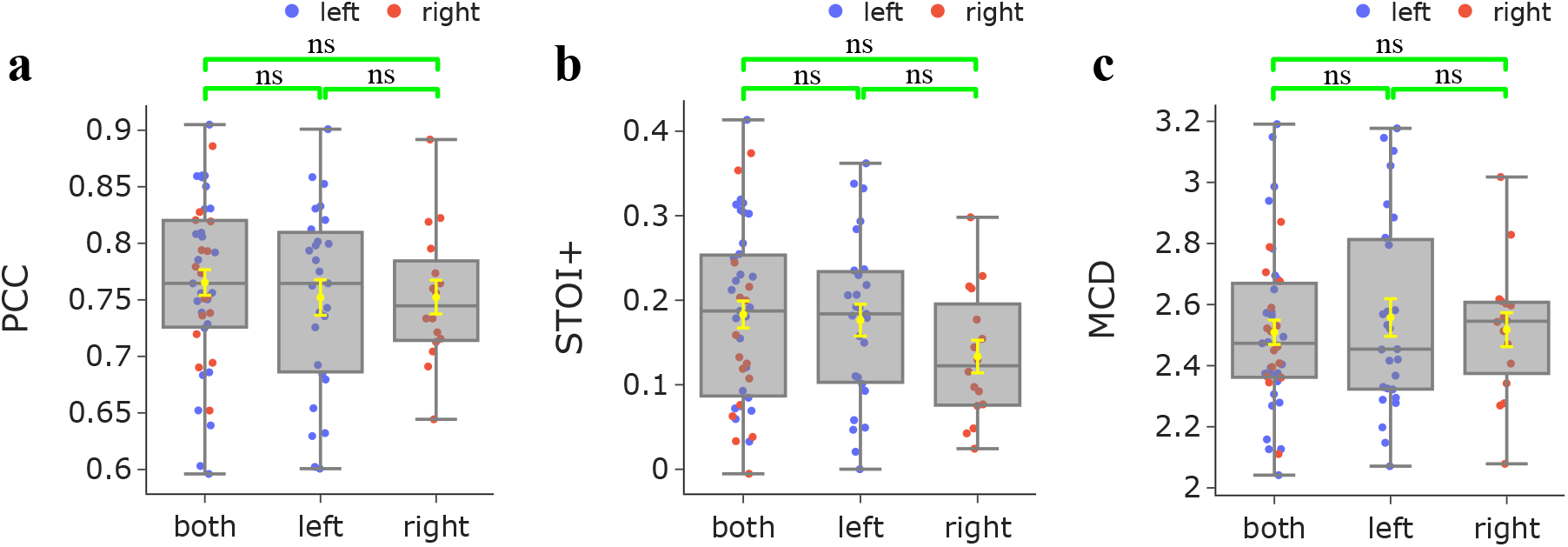
The comparison of speech decoding performance on unseen subjects between SwinTW trained on data from one hemisphere and SwinTW trained on data from both hemispheres. Models were trained separately for males and females. The results demonstrate that, compared with hemisphere-specific models, the SwinTW Neural Decoder trained on both hemispheres can achieve comparable or slightly better performance when inference on unseen subjects. Box plots as described in Fig. 3. Distributions were compared with each other as indicated. Green brackets indicate two experiments that are compared using the Wilcoxon rank-sum test. ns: P *>* 0.05.

## 4. Discussion

This study proposes a new Neural Decoder architecture, SwinTW, that does not have the grid-input assumption and can predict speech parameters from electrodes in any topological layout in the brain. The SwinTW removes the grid-based operations in the 3D Swin Transformer model used in our prior study [9] to make the model applicable for electrodes in any layout. Instead of relying on 2D grid indices to provide positional information about each electrode, the SwinTW relies on each electrode’s position in the standardized brain coordinate (i.e., MNI) and the brain region that the electrode resides in to generate relative positional bias for self-attention. The SwinTW was used as the Neural Decoder in the speech decoding pipeline proposed in our previous work [47, 9] and was trained using the 2-step training pipeline in [47, 9].

Our proposed SwinTW Neural Decoder achieved superior performance over the Neural Decoders based on ResNet and 3D Swin Transformer in [47, 9], which can only work with ECoG data. As illustrated in Figure 3, over 43 participants with low-density 8×8 ECoG electrodes, the SwinTW achieved higher mean PCC, STOI+, and lower MCD (PCC: 0.825, STOI+: 0.309, NCD: 2.341) than the ResNet (PCC:0.804, STOI+: 0.264, MCD: 2.374) and 3D Swin Transformer (PCC: mean: 0.785, STOI+: 0.216, MCD: 2.425) using the same 64 electrodes from the 8×8 ECoG grid. We attribute SwinTW’s better performance to its utilization of electrodes’ locations on the brain cortex (the MNI coordinate and brain region information) rather than the 2D grid index.

Unlike ResNet and 3D Swin Transformer, the SwinTW does not rely on 2D grid indices of electrodes and can accommodate both ECoG electrodes, strip and depth electrodes, and additional grid electrodes. Our results demonstrate that leveraging the additional electrodes can improve speech decoding performance, as illustrated in Figure 4. Specifically, for 39 subjects with additional active electrodes, the SwinTW utilizing the additional electrodes achieved better PCC (mean 0.838), STOI+ (mean 0.359), and MCD (mean 2.228) compared with the SwinTW using grid electrodes only (PCC: mean 0.825, STOI+: mean 0.318, MCD: mean 2.341). The superior results indicate that the neural activity recorded by the additional electrodes contains complementary information for decoding speech.

Our results further demonstrate that the SwinTW can achieve high decoding quality based only on sEEG electrodes. Specifically, as shown in Figure 4, for nine subjects with only sEEG electrodes implanted, we achieved PCC: mean 0.798, and STOI+: mean 0.341, and MCD: mean 2.396. The mean and range of PCC, STOI+, and MCD are slightly lower than the decoding performance obtained using ECoG electrodes but significantly higher than previously reported decoding performance from sEEG only, ranging between 0.54 to 0.77 in mean PCC [2, 3, 23, 45]. It is notable that there is no statistical difference between the decoding performance from sEEG vs. from ECoG or all electrodes in terms of STOI+ and MCD, the metrics that are better indicators of the intelligibility of the decoded speech. We present the distribution of electrode coverage in Figure S6, alongside the electrode contribution analysis in Figure S8, which quantifies the importance of each electrode across all participants. The methodology employed for the contribution analysis is detailed in [9]. The contribution analysis is conducted using subject-specific models. Despite the variation in electrode coverage across the grid, all-electrode, and sEEG cases, the contribution analysis reveals consistent patterns. Specifically, electrodes in the motor and temporal lobe regions exhibit greater contributions compared to those in other regions. Conversely, electrodes in other and unidentified regions show the lowest contributions, despite the relatively large number of electrodes present in these areas. This may explain why similar decoding performance can be achieved despite variations in electrode coverage. That is, the sEEG is sampling sufficient cortical regions in order to achieve similar decoding.

Since the SwinTW directly uses the anatomical positions of electrodes rather than their grid indices, it can be trained with data from multiple subjects. As shown in Figure 5, when evaluated on the testing trials of the participants in the training cohort and using only data on 8×8 grids, the resulting multi-subject model trained with data from 15 participants achieved statistically on par decoding performance (PCC: mean 0.837, STOI+: mean 0.352, MCD: mean 2.307), compared with the SwinTW trained for each subject individually (PCC: mean: 0.831, STOI+: mean: 0.335, and MCD: mean 2.313). This implies that the SwinTW model structure is able to effectively deal with the significant variability in the electrode placements among patients and make use of electrodes’ positions on the cortex. Previously, we have attempted to train ResNet and 3D Swin-based Neural Decoders using ECoG data from multiple participants. We were not able to improve the decoding performance compared to subject-specific models. That is likely because ResNet and 3D Swin models rely on the electrodes’ relative positions in the 2D grid. Because the ECoG grid is placed differently among the participants, the same relative difference in the 2D grid can be associated with very different anatomical positions in different participants, making using the grid index as positional information unsuitable when the data come from multiple participants.

Most significantly, the SwinTW model trained with multiple participants data demonstrated generalizability to participants outside the training cohorts, with high average decoding PCC (mean PCC = 0.765 over 43 unseen participants through a cross-validation study conducted separately for males and females). Figure 6 shows that the speech decoding performance achieved on unseen subjects overlaps significantly with that of the subject-specific model. Furthermore, a model trained with data from both the left and right hemispheres performs on par with those trained using only the left or right hemisphere on unseen participants (Figure 7). These results suggest that the SwinTW training using multiple participants’ data can successfully learn how to handle differences among subjects based on electrode signals and the anatomical position of the electrodes. The success of the left and right hemispheres co-training demonstrates the strong learning capacity of the SwinTW. The two-hemisphere co-training also allows the Neural Decoder to fully leverage the whole dataset as we no longer need to train the model separately for each hemisphere.

To summarize, the SwinTW Neural Decoder can predict speech parameters from electrode signals and electrodes’ positions on the brain cortex without requiring the electrodes to be arranged in a grid. The SwinTW Neural Decoder, in conjunction with our previously reported Speech Synthesizer, demonstrated superior speech decoding performance compared with our prior works based on ResNet and 3D Swin Transformers when only electrodes on a single ECoG array were used. Besides, the grid-free architecture of the SwinTW allows the model to leverage off-grid electrodes to improve speech decoding further. When using only sEEG data, the decoding performance was comparable with that using ECoG data. As explained in the Introduction, decoding speech from sEEG signals would have significant clinical advantages over using ECoG data. Furthermore, the SwinTW can be trained with data from multiple subjects regardless if the electrodes were implanted in the left or right brain hemispheres. The multi-subject SwinTW performed statistically on par with the subject-specific models for participants within the training cohort. Most importantly, our SwinTW trained with multiple participants’ data demonstrated good generalizability to subjects outside the training cohorts, achieving high average decoding PCC.

We are one of the few studies ([42, 11, 31]) demonstrating speech-decoding models trained across multiple participants. However, these other prior works embed subject-specific layers in their model structures, and hence, the models need subject-specific data for training. To our knowledge, we are the first to design a framework that goes beyond subject-specific training without using subject-specific layers. Our result demonstrates the exciting possibility of developing speech prostheses without collecting subject-specific training data: We can train a reliable decoder with data from selected participants and then directly deploy the model to a new participant. Note that although our experiments on the multi-subject model only considered the ECoG grid data, we expect similar trends when using ECoG plus non-grid data or non-grid data only.

Notably, the proposed SwinTW Neural Decoder is not limited to being used with our speech synthesizer. It could potentially be used to decode other latent features, e.g. the HuBERT latent features [18], which can then drive a corresponding synthesizer [26]. The work in [32] successfully decoded speech with high word decoding accuracy by decoding to quantized HuBERT units using an RNN decoder from high-density ECoG signals of a single participant. However, the RNN structure cannot be trained with multi-subject data without introducing subject-specific layers. It will be interesting to explore the potential of training a SwinTW decoder using data from multiple participants with surface and/or depth electrodes, to map the neural signals to the HuBERT units and compare the decoding performance with the subject-specific RNN model or multi-subject RNN models with subject-specific layers.

One limitation of our current study is that the decoding performance for participants outside of the training cohorts is not consistently high. This could be potentially solved by including more participants in the training set when larger datasets become available. Furthermore, the SwinTW model structure can also be extended to include subject-specific layers for improved performance. We would explore training the non-subject-specific layers with a large pre-collected multi-subject dataset and refine only the subject-specific layer with a small amount of data for any new participants.

## Data availability

The data of this study are available from the corresponding author upon request. Although all participants consented to share their data for research purposes, not all participants agreed to share their audio publicly. Given the sensitive nature of audio speech data, we will share data with researchers that directly contact the corresponding author and provide documentation that the data will be strictly used for research purposes and will comply with the terms of our study NYU Langone IRB.

## Acknowledgments

This material is based upon work supported by the National Science Foundation under Grant No. IIS-1912286 and IIS-2309057 (Y.W., A.F.) and National Institute of Health R01NS109367, R01NS115929, R01DC018805 (A.F.).

## References

[1] M. Angrick, C. Herff, E. Mugler, M. C. Tate, M. W. Slutzky, D. J. Krusienski, and T. Schultz. Speech synthesis from ecog using densely connected 3d convolutional neural networks. Journal of neural engineering, 16(3):036019, 2019.

[2] M. Angrick, M. Ottenhoff, L. Diener, D. Ivucic, G. Ivucic, S. Goulis, A. J. Colon, L. Wagner, D. J. Krusienski, P. L. Kubben, et al. Towards closed-loop speech synthesis from stereotactic eeg: a unit selection approach. In ICASSP 2022-2022 IEEE International Conference on Acoustics, Speech and Signal Processing (ICASSP), pages 1296–1300. IEEE, 2022.

[3] M. Angrick, M. Ottenhoff, L. Diener, D. Ivucic, G. Ivucic, S. Goulis, J. Saal, A. Colon, L. Wagner, D. Krusienski, et al. Real-time synthesis of imagined speech processes from minimally invasive recordings of neural activity. commun biol 4 (1): 1055–1055, 2021.

[4] G. K. Anumanchipalli, J. Chartier, and E. F. Chang. Speech synthesis from neural decoding of spoken sentences. Nature, 568(7753):493–498, 2019.

[5] J. L. Ba, J. R. Kiros, and G. E. Hinton. Layer normalization. arXiv preprint arXiv:1607.06450, 2016.

[6] P. Boersma and V. Van Heuven. Speak and unspeak with praat. Glot International, 5(9/10):341–347, 2001.

[7] J. S. Brumberg, A. Nieto-Castanon, P. R. Kennedy, and F. H. Guenther. Brain–computer interfaces for speech communication. Speech communication, 52(4):367–379, 2010.

[8] S. Chakrabarti, H. M. Sandberg, J. S. Brumberg, and D. J. Krusienski. Progress in speech decoding from the electrocorticogram. Biomedical Engineering Letters, 5:10–21, 2015.

[9] X. Chen, R. Wang, A. Khalilian-Gourtani, L. Yu, P. Dugan, D. Friedman, W. Doyle, O. Devinsky, Y. Wang, and A. Flinker. A neural speech decoding framework leveraging deep learning and speech synthesis. Nature Machine Intelligence, pages 1–14, 2024.

[10] B. C. Chiluba. Tackling disability of speech due to stroke: Perspectives from stroke caregivers of the university teaching hospital in zambia. Indonesian Journal of Disability Studies, 6(2):215–222, 2019.

[11] A. Défossez, C. Caucheteux, J. Rapin, O. Kabeli, and J.-R. King. Decoding speech perception from non-invasive brain recordings. Nature Machine Intelligence, 5(10):1097–1107, 2023.

[12] A. Dosovitskiy, L. Beyer, A. Kolesnikov, D. Weissenborn, X. Zhai, T. Unterthiner, M. Dehghani, M. Minderer, G. Heigold, S. Gelly, et al. An image is worth 16×16 words: Transformers for image recognition at scale. arXiv preprint arXiv:2010.11929, 2020.

[13] J. Engel, L. Hantrakul, C. Gu, and A. Roberts. Ddsp: Differentiable digital signal processing. arXiv preprint arXiv:2001.04643, 2020.

[14] S. Graetzer and C. Hopkins. Intelligibility prediction for speech mixed with white gaussian noise at low signal-to-noise ratios. The Journal of the Acoustical Society of America, 149(2):1346–1362, 2021.

[15] K. He, X. Zhang, S. Ren, and J. Sun. Deep residual learning for image recognition. In Proceedings of the IEEE conference on computer vision and pattern recognition, pages 770–778, 2016.

[16] C. Herff, L. Diener, M. Angrick, E. Mugler, M. C. Tate, M. A. Goldrick, D. J. Krusienski, M. W. Slutzky, and T. Schultz. Generating natural, intelligible speech from brain activity in motor, premotor, and inferior frontal cortices. Frontiers in neuroscience, 13:1267, 2019.

[17] C. Herff, D. J. Krusienski, and P. Kubben. The potential of stereotactic-eeg for brain-computer interfaces: current progress and future directions. Frontiers in neuroscience, 14:123, 2020.

[18] W.-N. Hsu, B. Bolte, Y.-H. H. Tsai, K. Lakhotia, R. Salakhutdinov, and A. Mohamed. Hubert: Self-supervised speech representation learning by masked prediction of hidden units. IEEE/ACM Transactions on Audio, Speech, and Language Processing, 29:3451–3460, 2021.

[19] K. Iida and H. Otsubo. Stereoelectroencephalography: indication and efficacy. Neurologia medico-chirurgica, 57(8):375–385, 2017.

[20] M. Jacobs and C. Ellis. Aphasianomics: estimating the economic burden of poststroke aphasia in the united states. Aphasiology, 37(1):25–38, 2023.

[21] A. Khalilian-Gourtani, R. Wang, X. Chen, L. Yu, P. Dugan, D. Friedman, W. Doyle, O. Devinsky, Y. Wang, and A. Flinker. A corollary discharge circuit in human speech. BioRxiv, pages 2022–09, 2022.

[22] D. P. Kingma and J. Ba. Adam: A method for stochastic optimization. arXiv preprint arXiv:1412.6980, 2014.

[23] J. Kohler, M. C. Ottenhoff, S. Goulis, M. Angrick, A. J. Colon, L. Wagner, S. Tousseyn, P. L. Kubben, and C. Herff. Synthesizing speech from intracranial depth electrodes using an encoder-decoder framework. arXiv preprint arXiv:2111.01457, 2021.

[24] S. Komeiji, K. Shigemi, T. Mitsuhashi, Y. Iimura, H. Suzuki, H. Sugano, K. Shinoda, and T. Tanaka. Transformer-based estimation of spoken sentences using electrocorticography. In ICASSP 2022-2022 IEEE International Conference on Acoustics, Speech and Signal Processing (ICASSP), pages 1311–1315. IEEE, 2022.

[25] J. Kominek, T. Schultz, and A. W. Black. Synthesizer voice quality of new languages calibrated with mean mel cepstral distortion. In SLTU, pages 63–68, 2008.

[26] K. Lakhotia, E. Kharitonov, W.-N. Hsu, Y. Adi, A. Polyak, B. Bolte, T.-A. Nguyen, J. Copet, A. Baevski, A. Mohamed, et al. On generative spoken language modeling from raw audio. Transactions of the Association for Computational Linguistics, 9:1336–1354, 2021.

[27] Z. Liu, H. Hu, Y. Lin, Z. Yao, Z. Xie, Y. Wei, J. Ning, Y. Cao, Z. Zhang, L. Dong, et al. Swin transformer v2: Scaling up capacity and resolution. In Proceedings of the IEEE/CVF conference on computer vision and pattern recognition, pages 12009–12019, 2022.

[28] Z. Liu, Y. Lin, Y. Cao, H. Hu, Y. Wei, Z. Zhang, S. Lin, and B. Guo. Swin transformer: Hierarchical vision transformer using shifted windows. In Proceedings of the IEEE/CVF international conference on computer vision, pages 10012–10022, 2021.

[29] Z. Liu, J. Ning, Y. Cao, Y. Wei, Z. Zhang, S. Lin, and H. Hu. Video swin transformer. In Proceedings of the IEEE/CVF conference on computer vision and pattern recognition, pages 3202–3211, 2022.

[30] S. Luo, Q. Rabbani, and N. E. Crone. Brain-computer interface: applications to speech decoding and synthesis to augment communication. Neurotherapeutics, 19(1):263–273, 2022.

[31] J. G. Makin, D. A. Moses, and E. F. Chang. Machine translation of cortical activity to text with an encoder–decoder framework. Nature neuroscience, 23(4):575–582, 2020.

[32] S. L. Metzger, K. T. Littlejohn, A. B. Silva, D. A. Moses, M. P. Seaton, R. Wang, M. E. Dougherty, J. R. Liu, P. Wu, M. A. Berger, et al. A high-performance neuroprosthesis for speech decoding and avatar control. Nature, pages 1–10, 2023.

[33] D. A. Moses, M. K. Leonard, J. G. Makin, and E. F. Chang. Real-time decoding of question-and-answer speech dialogue using human cortical activity. Nature communications, 10(1):3096, 2019.

[34] D. A. Moses, S. L. Metzger, J. R. Liu, G. K. Anumanchipalli, J. G. Makin, P. F. Sun, J. Chartier, M. E. Dougherty, P. M. Liu, G. M. Abrams, A. T.-C. D.O., K. Ganguly, and E. F. Chang. Neuroprosthesis for decoding speech in a paralyzed person with anarthria. New England Journal of Medicine, 385(3):217–227, 2021.

[35] L. E. Nicholas and R. H. Brookshire. Comprehension of spoken narrative discourse by adults with aphasia, right-hemisphere brain damage, or traumatic brain injury. American Journal of Speech-Language Pathology, 4(3):69–81, 1995.

[36] N. F. Ramsey, E. Salari, E. J. Aarnoutse, M. J. Vansteensel, M. G. Bleichner, and Z. Freudenburg. Decoding spoken phonemes from sensorimotor cortex with high-density ecog grids. Neuroimage, 180:301–311, 2018.

[37] R. W. Schafer. What is a savitzky-golay filter?[lecture notes]. IEEE Signal processing magazine, 28(4):111–117, 2011.

[38] T. Schultz, M. Wand, T. Hueber, D. J. Krusienski, C. Herff, and J. S. Brumberg. Biosignal-based spoken communication: A survey. IEEE/ACM Transactions on Audio, Speech, and Language Processing, 25(12):2257–2271, 2017.

[39] J. Senda, M. Tanaka, K. Iijima, M. Sugino, F. Mori, Y. Jimbo, M. Iwasaki, and K. Kotani. Auditory stimulus reconstruction from ecog with dnn and self-attention modules. Biomedical Signal Processing and Control, 89:105761, 2024.

[40] K. Shigemi, S. Komeiji, T. Mitsuhashi, Y. Iimura, H. Suzuki, H. Sugano, K. Shinoda, K. Yatabe, and T. Tanaka. Synthesizing speech from ecog with a combination of transformer-based encoder and neural vocoder. In ICASSP 2023-2023 IEEE International Conference on Acoustics, Speech and Signal Processing (ICASSP), pages 1–5. IEEE, 2023.

[41] J. Shum, L. Fanda, P. Dugan, W. K. Doyle, O. Devinsky, and A. Flinker. Neural correlates of sign language production revealed by electrocorticography. Neurology, 95(21):e2880–e2889, 2020.

[42] P. Sun, G. K. Anumanchipalli, and E. F. Chang. Brain2char: a deep architecture for decoding text from brain recordings. Journal of neural engineering, 17(6):066015, 2020.

[43] N. Tandon, B. A. Tong, E. R. Friedman, J. A. Johnson, G. Von Allmen, M. S. Thomas, O. A. Hope, G. P. Kalamangalam, J. D. Slater, and S. A. Thompson. Analysis of morbidity and outcomes associated with use of subdural grids vs stereoelectroencephalography in patients with intractable epilepsy. JAMA neurology, 76(6):672–681, 2019.

[44] R. Thomas, A. M. O’Connor, and S. Ashley. Speech and language disorders in patients with high grade glioma and its influence on prognosis. Journal of neuro-oncology, 23:265–270, 1995.

[45] M. Verwoert, M. C. Ottenhoff, S. Goulis, A. J. Colon, L. Wagner, S. Tousseyn, J. P. van Dijk, P. L. Kubben, and C. Herff. Dataset of speech production in intracranial electroencephalography. Scientific data, 9(1):434, 2022.

[46] R. Wang, X. Chen, A. Khalilian-Gourtani, Z. Chen, L. Yu, A. Flinker, and Y. Wang. Stimulus speech decoding from human cortex with generative adversarial network transfer learning. In 2020 IEEE 17th International Symposium on Biomedical Imaging (ISBI), pages 390–394. IEEE, 2020.

[47] R. Wang, X. Chen, A. Khalilian-Gourtani, L. Yu, P. Dugan, D. Friedman, W. Doyle, O. Devinsky, Y. Wang, and A. Flinker. Distributed feedforward and feedback cortical processing supports human speech production. Proceedings of the National Academy of Sciences, 120(42):e2300255120, 2023.

[48] X. Wu, S. Wellington, Z. Fu, and D. Zhang. Speech decoding from stereo-electroencephalography (seeg) signals using advanced deep learning methods. Journal of Neural Engineering, 2024.

