## supplementary file for "Subject-Agnostic Transformer-Based Neural Speech Decoding from Surface and Depth Electrode Signals"

### CONTENTS

1 Random Shuffled Model Results 2

2 Electrode Coverage Distribution 4

3 Additional Electrodes 6

4 Electrodes Contribution 7

5 Supplementary Video Demo 9

References 10

### 1. RANDOM SHUFFLED MODEL RESULTS

To assess the chance accuracy, we randomly shuffled the input ECoG and sEEG signals temporally and fed them into the model trained with correctly aligned data. Then we compute the metrics for such shuffled input. We replaced all the decoding performance plots by adding the decoding performance with shuffled data (Fig. S1, S2, S3, S4). It is evident that the decoding performance with non-shuffled data is significantly better.

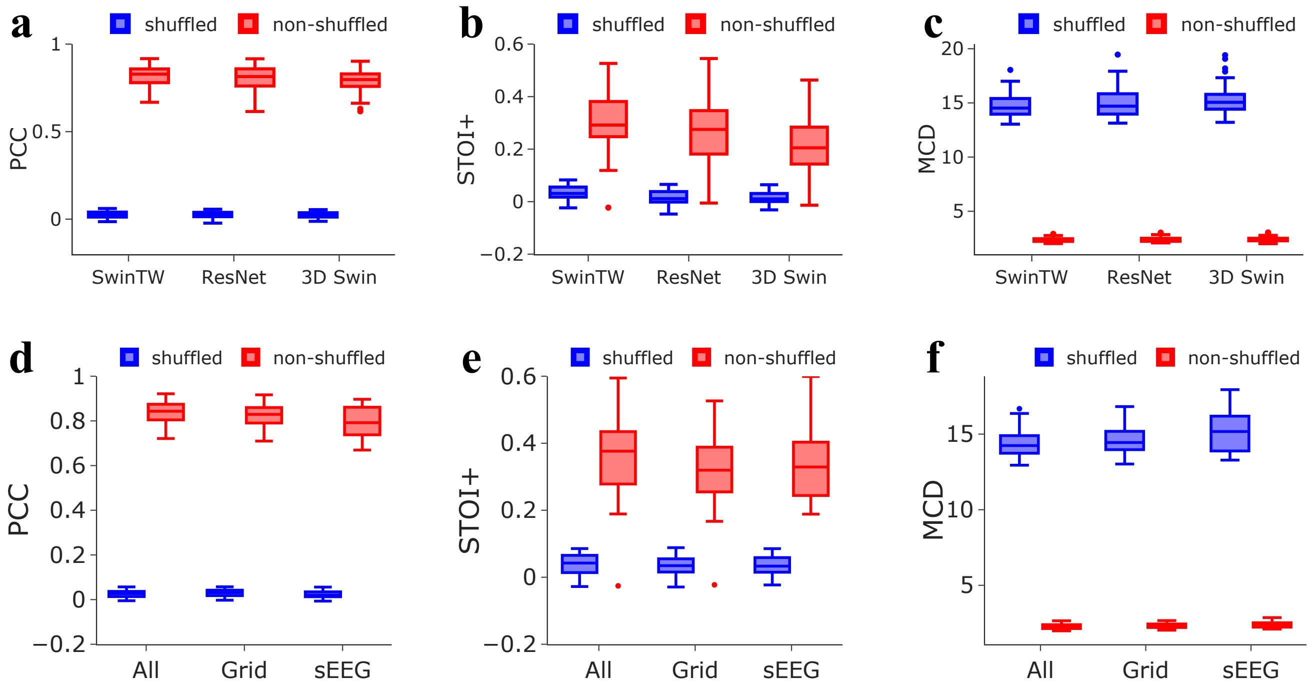


**Fig. S1.** | Supplementary Fig S1.

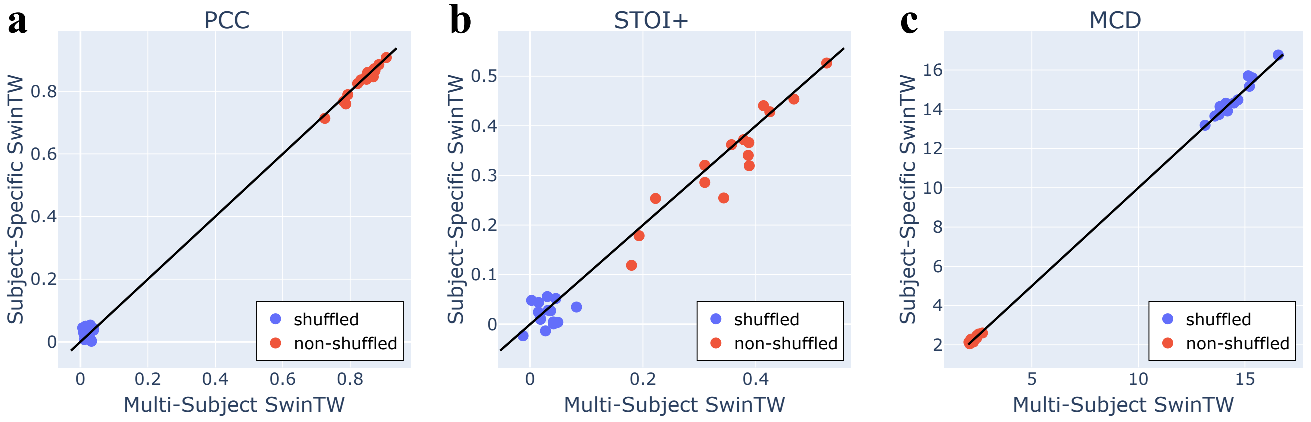


**Fig. S2.** | Supplementary Fig S2.

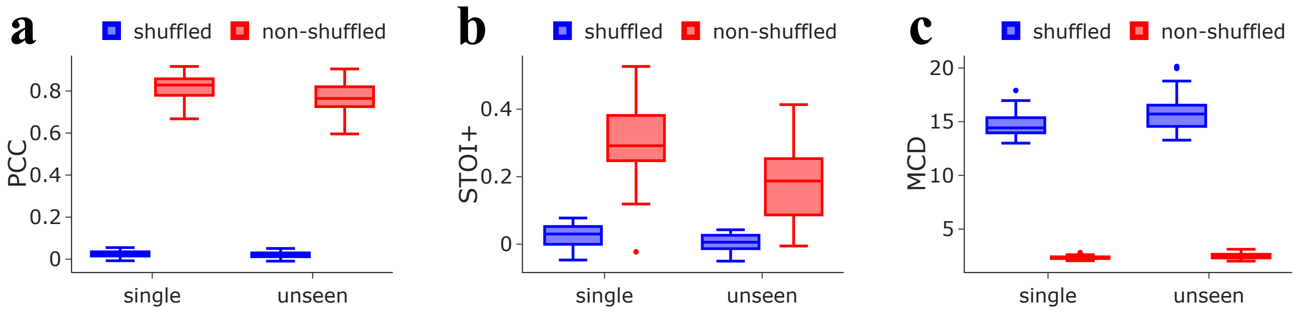


**Fig. S3.** | Supplementary Fig S3.

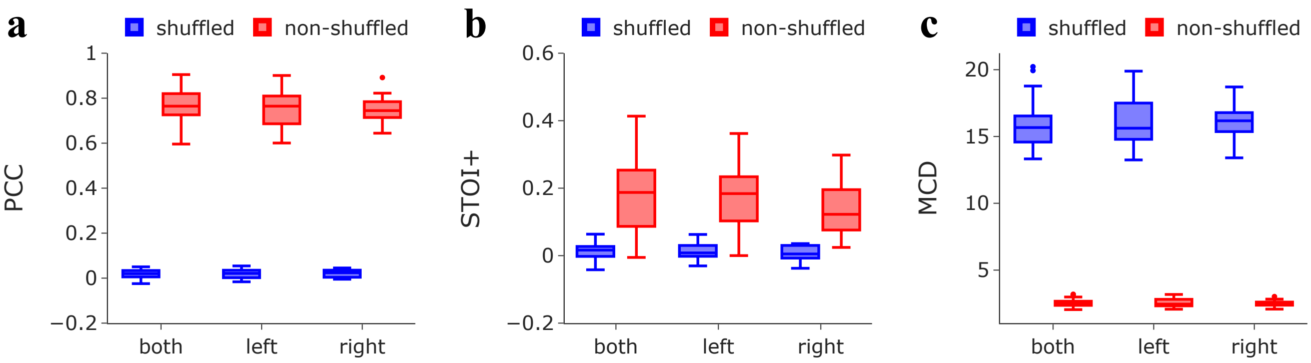


**Fig. S4.** | Supplementary Fig S4.

### 2. ELECTRODE COVERAGE DISTRIBUTION

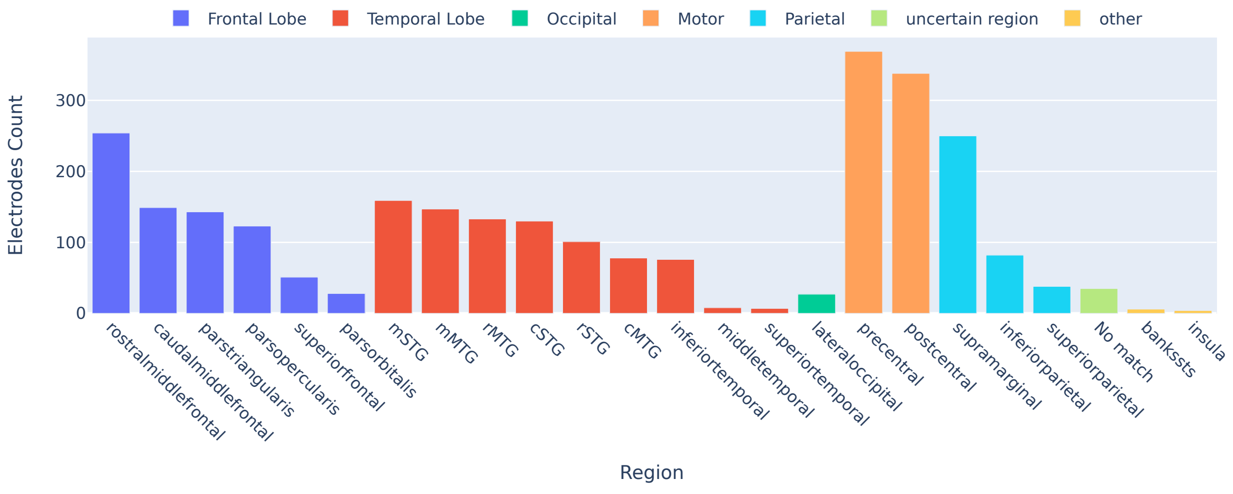


**(a)** Grid electrodes distribution over the brain across 43 Participants with ECoG electrodes

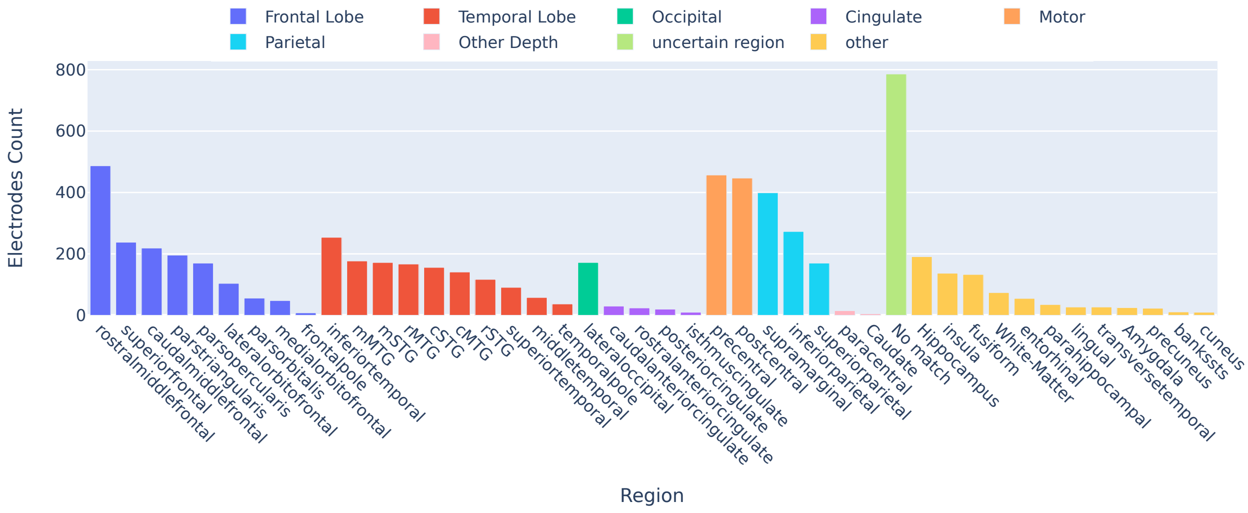


**(b)** All electrodes distribution over the brain across 43 Participants with ECoG and other electrodes

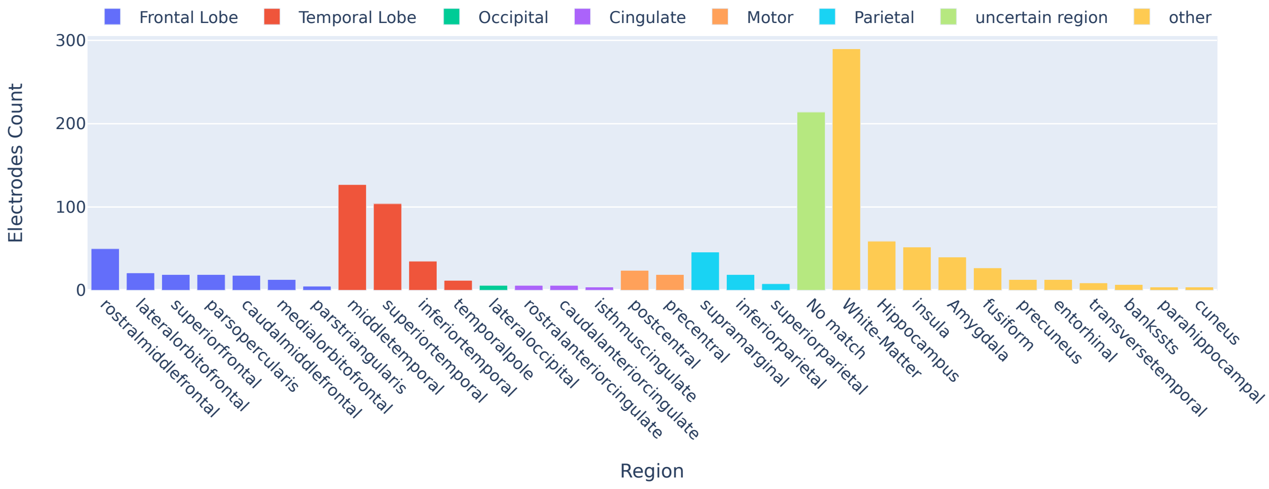


**(c)** Depth electrodes distribution over the brain across 9 Participants with sEEG electrodes only

**Fig. S6.** Electrodes Distribution

### 3. ADDITIONAL ELECTRODES

The proposed SwinTW can easily leverage off-grid electrodes (blue electrodes in Fig. S7) to provide additional information for speech decoding. In our study, for each participant with additional electrodes beyond one ECoG grid, we selected additional electrodes with a standard deviation of the signal greater than a threshold, determined following the approach described in [1] for identifying active electrodes.

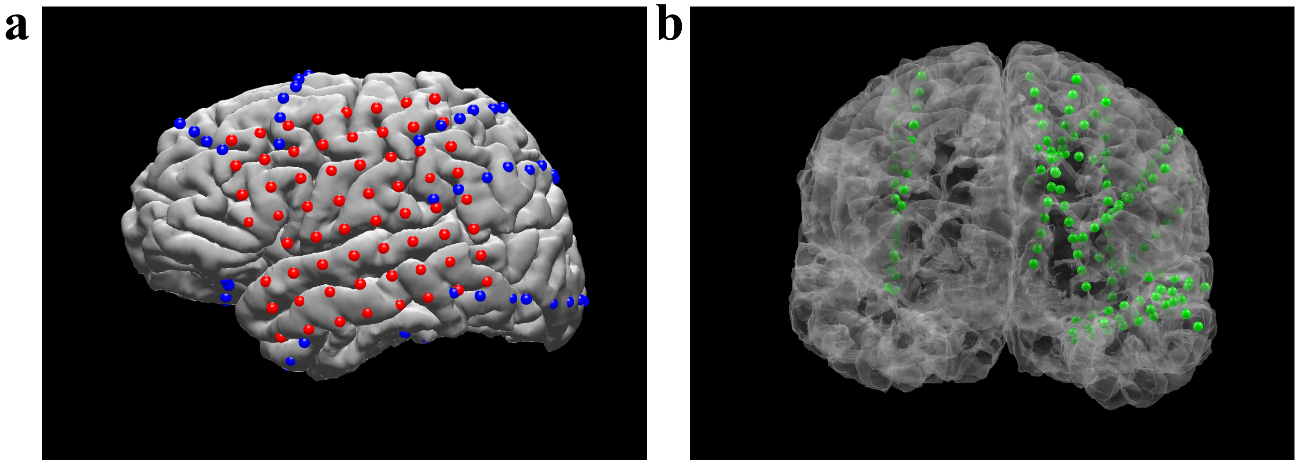


**Fig. S7.** | Illustration of non-gird electrodes used. Blue indicates strip electrodes, and green indicates depth electrodes.

### 4. ELECTRODES CONTRIBUTION

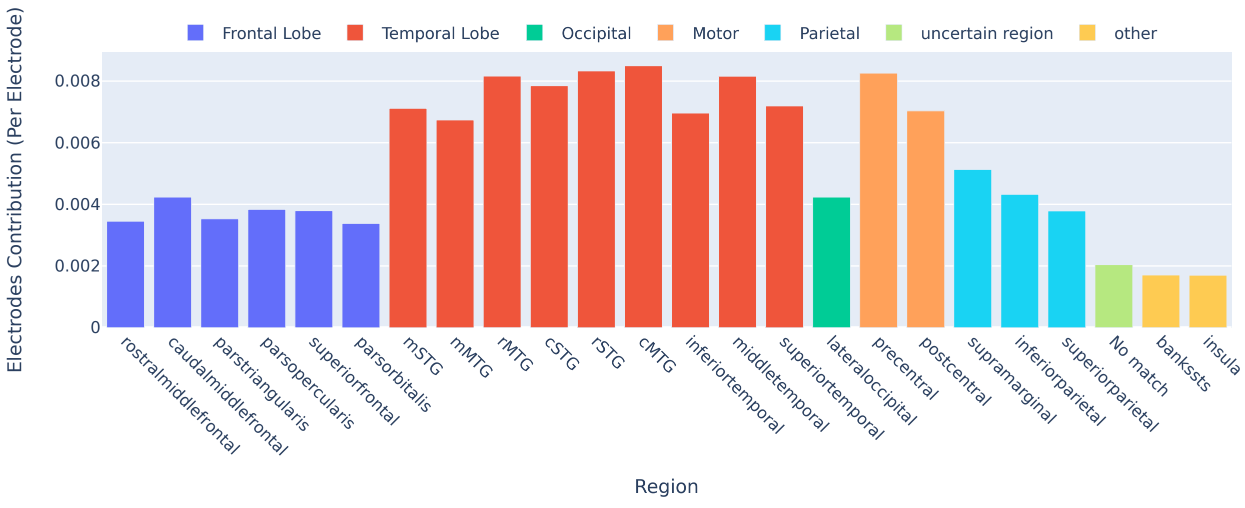


**(a)** Grid electrodes contribution averaged over 43 Participants

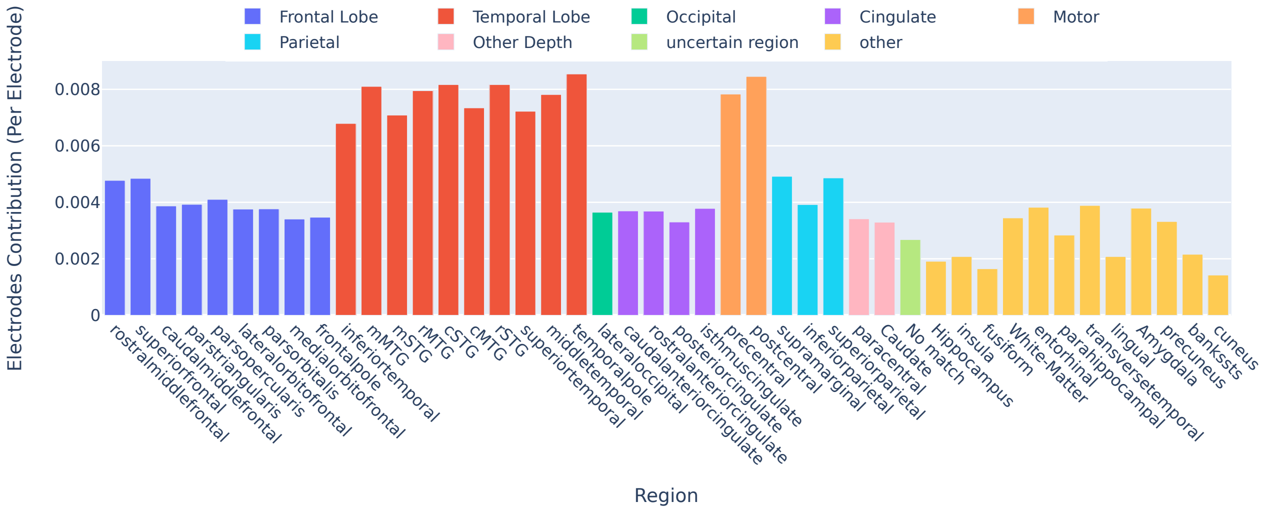


**(b)** All electrodes contribution averaged over 39 Participants

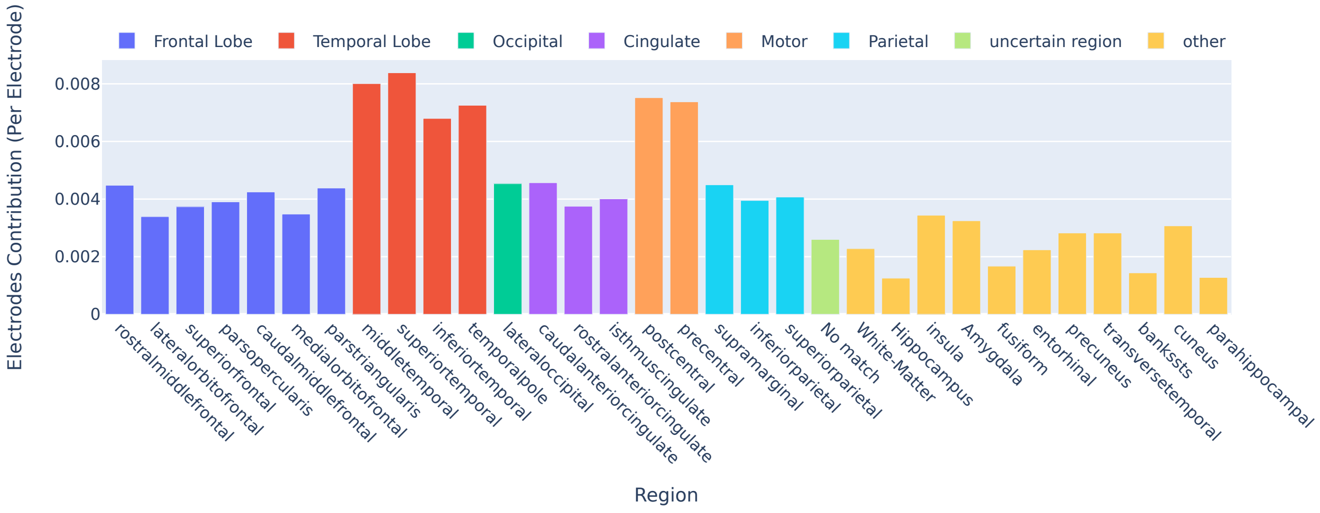


**(c)** Depth electrodes contribution averaged over 9 sEEG Participants

**Fig. S8.** Electrodes Distribution

### 5. SUPPLEMENTARY VIDEO DEMO

We have added the video containing sample decoded speech and linked it here: https://xc1490.github.io/swinTW

### REFERENCES

1. A. Khalilian-Gourtani, R. Wang, X. Chen, L. Yu, P. Dugan, D. Friedman, W. Doyle, O. Devinsky, Y. Wang, and A. Flinker, "A corollary discharge circuit in human speech," BioRxiv pp. 2022–09 (2022).
